# Engineering a Functional small RNA Negative Autoregulation Network with Model-guided Design

**DOI:** 10.1101/227637

**Authors:** Chelsea Y. Hu, Melissa K. Takahashi, Yan Zhang, Julius B. Lucks

## Abstract

RNA regulators are powerful components of the synthetic biology toolbox. Here, we expand the repertoire of synthetic gene networks built from these regulators by constructing a transcriptional negative autoregulation (NAR) network out of small RNAs (sRNAs). NAR network motifs are core motifs of natural genetic networks, and are known for reducing network response time and steady state signal noise. Here we use cell-free transcription-translation (TX-TL) reactions and a computational model to design and prototype sRNA NAR constructs. Using parameter sensitivity analysis, we design a simple set of experiments that allow us to accurately predict NAR function in TX-TL. We transfer successful network designs *in vivo* and show that our sRNA transcriptional network reduces both network response time and noise in steady-state gene expression. This work broadens our ability to construct increasingly sophisticated RNA genetic networks with predictable function.

## Introduction

A major goal of synthetic biology is to design and construct synthetic gene networks that can be used to reprogram the behavior of living cells to accomplish specialized functions. RNA regulators have emerged as powerful tools in the synthetic biology toolbox of gene expression control that can be used to achieve this goal^1^. Specifically, synthetic biologists have engineered a diverse array of RNA regulators that can be used to control many of the core processes of gene expression including transcription^2–5^, translation^6–8^ and mRNA degradation^9–12^. In addition to being able to tune the expression of individual genes, these engineered RNA regulators have also been connected together to create synthetic gene networks in the form of logic gates^2,3,5,7,13^, cascades^2,5,13–17^, single input modules^5,16^ feed forward loops^18^, and hybrid feedback controllers^12^. While these synthetic network motifs showcase some of the advantages of RNA regulators, they represent the beginning of larger efforts to create a greater diversity of RNA network function that is necessary to engineer higher order cellular behavior.

Much of the design of new synthetic genetic networks has been guided by the understanding and repurposing of natural genetic networks that control a large range of cellular functions including cellular maintenance, development, and the response to environmental perturbations. These natural genetic networks are composed of repeating occurrences of simple patterns called network motifs^19^. Examples of network motifs include autoregulation where regulators can regulate their own synthesis, feedback loops where regulators can regulate upstream portions of a network, and feed-forward loops where regulators can regulate downstream portions of a network, among others^19^. Each of these network motifs has a distinct information processing function, and larger networks are made by composing motifs in different ways to achieve higher order information processing. In essence, these network motifs form a basis set of elements from which larger networks can be built, making the construction and characterization of synthetic versions a powerful target for synthetic biology^19^.

The simplest motif in transcription networks is negative autoregulation (NAR)^19^, a motif in which transcription factors negatively regulate their own transcription^20^. One of the key functions of NARs in nature is to speed the response time of a transcription network^21^. NARs are also known to reduce fluctuations, or “noise”, of steady state gene expression of genetic circuitry^22–24^. NARs are also a key element of genetic oscillator modules^25^ as well as biological feedback controllers^26^. Thus NARs represent a rich target to construct using RNA genetic circuitry, which may have particular advantages due to the fast dynamics observed for RNA mediated networks^27^.

In this work, we engineer the first sRNA transcriptional NAR network using the transcriptional attenuator from the *Staphylococcus aureus* plasmid pT181^28^. To do this, we leverage recent advances in using cell-free transcription-translation (TX-TL) reactions^16^ and a computational modeling framework^29^ to rapidly prototype and characterize NAR constructs that are then transferred to living cells. We first simulate RNA-based NAR networks qualitatively with a simplified model to anticipate challenges of building an RNA-based NAR network. Following the model’s suggestion that increased copies of the RNA repressor would be needed for observable NAR function, we construct and optimize new RNA parts to construct the NAR network. We then develop a quantitative model of the network by representing the biological processes as a system of ordinary differential equations (ODEs) and use a sensitivity analysis approach to design parameterization experiments. Next, we estimate all unknown parameters of the NAR network and predict its dynamic trajectories in TX-TL reactions quantitatively. Comparison of the simulation against experimental data shows that the simulation captures the experimental result within a 95% confidence interval. Moreover, we show that the NAR network speeds up the response time for both simulated and experimental dynamic trajectories. Simulations also suggest that our sRNA transcriptional NAR reduces an aspect of the noise of the steady state signal. Finally, we move the TX-TL-prototyped NAR network into *E. coli* where we show that our NAR motif also significantly reduces network response time as well as the noise of the steady state signal *in vivo*.

## Results and discussion

### Using simplified computational models to guide NAR design

To build an RNA-based NAR network, we sought to use the pT181 transcriptional attenuator^28^ which we have previously used to build RNA-only transcriptional networks^2,3,27^. The pT181 attenuator is an RNA sequence in the 5’ untranslated region of a gene that regulates transcription elongation through the formation of an intrinsic terminator hairpin (Figure 1A). In its native or ON state, intramolecular interactions with an anti-terminator sequence prevents terminator formation and allows transcription elongation. In its OFF state, interactions with a repressor sRNA, which is complementary to the upstream region of the attenuator, sequesters the anti-terminator sequence, allowing the terminator hairpin to form and preventing transcription elongation of the downstream gene (Figure S1A)^28,30^. Thus the combination of repressor sRNA and attenuator act as an RNA-based transcriptional repressor.

**Figure 1.**
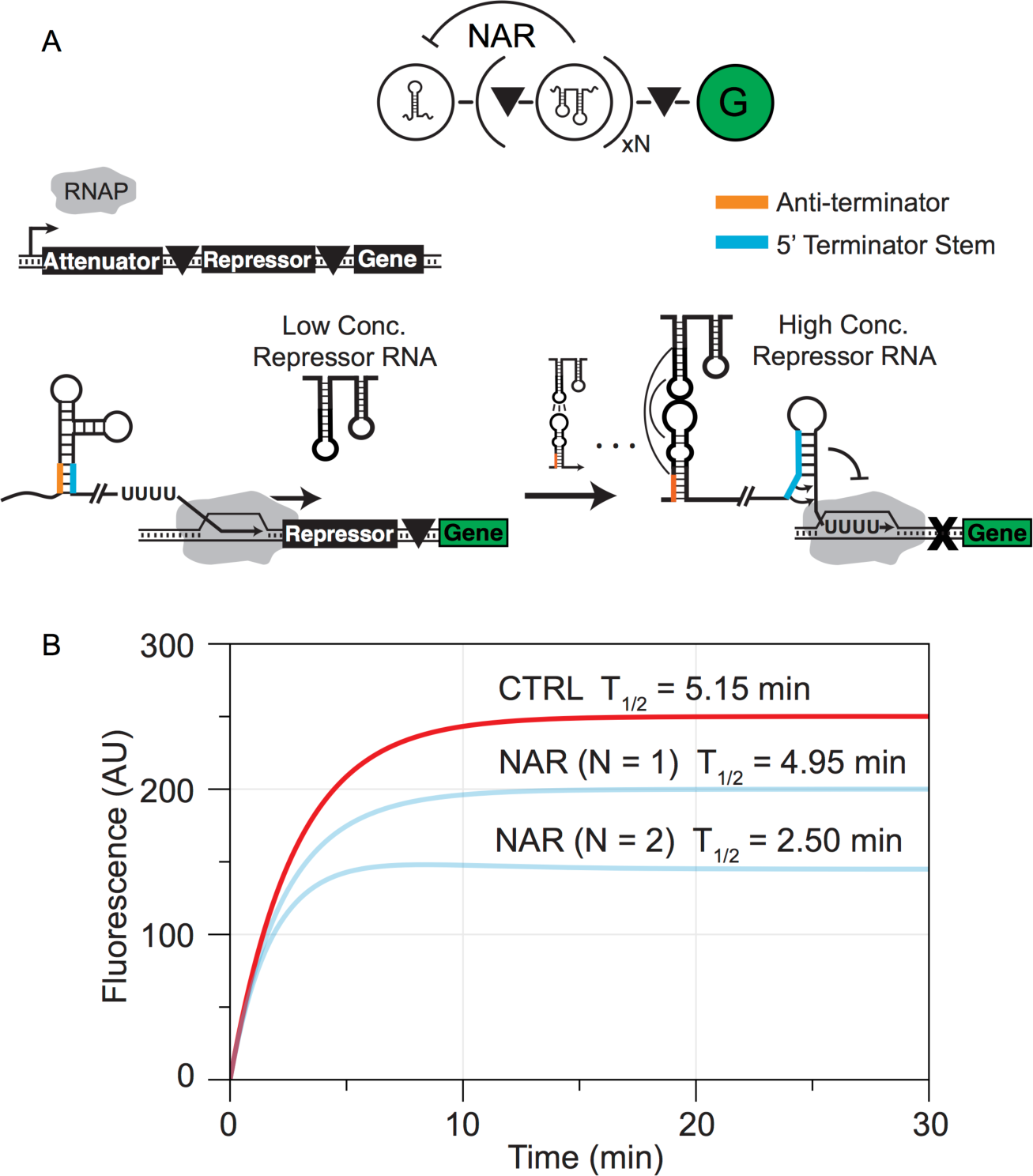
A simple model of the proposed sRNA transcriptional NAR construct uncovers potential challenges and provides design guidelines for network construction. (A) The transcriptional NAR network motif and proposed sRNA implementation. The pT181 attenuator regulates the transcription of N copies of its own repressor RNA (repressor, R) and a reporter gene (G), each insulated from each other by self-cleaving ribozymes (triangles). Once transcription is activated, initially RNA polymerase is allowed to transcribe the repressor RNA-reporter gene construct. As concentration of repressor RNA increases, it binds to the attenuator, which leads to intrinsic terminator formation and repression of transcription of the repressor RNA-reporter gene construct. (B) A simplified two-component model that describes the synthesis and degradation of the repressor (R) and reporter gene (G) provides insight for network construction. The dynamic trajectories were simulated by this model with parameters adapted from a previous study (Table S1)^29^. The simulation suggests that a single copy of repressor RNA (N=1) would be insufficient to significantly reduce network response time, while tandem copies of repressor RNA (N=2) would.

To build an NAR network, we first proposed a construct in which the pT181 attenuator regulates the transcription of its own repressor sRNA, followed by a reporter gene (Figure 1A). At the start, the attenuator would allow transcription of both the repressor sRNA and reporter gene until the concentration of repressor sRNA builds up and binds to the attenuator. Interaction of the repressor sRNA with the attenuator would lead to terminator formation and prevent further transcription of repressor sRNA and reporter gene.

We first sought to determine the feasibility of this pT 181 attenuator based NAR network design. To do this, we developed a simple mathematical model of the network using parameters from previous work where we simulated a transcriptional cascade in TX-TL using the pT181 attenuator^29^. Since the RNA-based NAR would uniquely operate at the RNA level, we sought a reporter that would allow us to directly measure and model RNA levels within the system. We therefore proposed an NAR construct that used the malachite green (MG)-binding RNA aptamer as a reporter^31^. Once folded properly, the RNA aptamer can bind the small molecule chromophore malachite green (MG), which results in a fluorescence output. The MG aptamer therefore allows experimental characterization of the RNA levels of the network with a convenient fluorescence read out^31^, and would allow us to directly model time trajectories of our experimental observable.

Our initial simplified model treated the basic gene expression processes of the NAR network as coupled chemical reactions of two species: the repressor sRNA (R) and the reporter malachite green RNA (Figure 1B). We considered the concentrations of these two species in time to be functions of their constant synthesis and degradation rates. For simplicity, we modeled the transcriptional repression with a first order Hill function based on previous experimental work^2^, in which K represents the concentration of active repressor that is required to achieve significant repression of downstream gene transcription (Figure 1B). Finally, to simulate the model, we used estimated parameter values based on the parameters we obtained in previous work in modeling RNA-based transcriptional cascades in TXTL reactions^29^ (Table S1).

One of the defining characteristics of an NAR network is that it speeds up a network’s response time, defined as the time it takes for a dynamic process to reach half of its steady state level^21^. We therefore used response time as an indicator of a functional NAR network. The initial simulation of the NAR network showed an insignificant decrease in response time compared to a control network where the negative feedback is broken (4.95 min compared to 5.15 min) (Figure 1B). We hypothesized this was due to the fast degradation rate of repressor sRNA and weak repression of the attenuator that prohibits sufficient accumulation of repressor sRNA to achieve a significant repression. To solve this potential problem, we proposed a new NAR design that increased the concentration of repressor sRNA by including two copies of the RNA in tandem. For convenience, in the simple model we treated this configuration as a new repressor sRNA species that is twice as repressive, but that also has a degradation rate which is half of that of a single repressor RNA to account for its increase in size. The simulation of this construct suggested that by using two copies of repressor sRNA, the response time would decrease from T_1/2_=5.15 minutes to T_1/2_= 2.50 minutes (Figure 1B).

Our two-dimensional model thus allowed us to capture the essence of an RNA-based NAR network and predict qualitative response times with minimal investigation. Based on the guidance of this simplified model, we next sought to implement an RNA-based NAR network with tandem copies of repressor sRNAs.

### Optimization of RNA parts for NAR construction

Overall, the proposed NAR architecture consists of four distinct RNA parts that must be fused together in tandem on the same RNA molecule for proper function: an upstream attenuator to control transcription, two copies of the repressor sRNA to provide the negative feedback if synthesized, and the MG aptamer to provide a fluorescent read-out (Figure 1). In addition, there are accessory transcriptional terminators on the ends of constructs that are needed to define expression constructs. A functional NAR requires each individual RNA part to fold into proper secondary structures, which is increasingly difficult with more RNA parts in tandem. To help alleviate potential problems with mis-folding and mis-function, we therefore sought to minimize the size of the parts involved in the construct. Specifically, we sought to minimize the length of the repressor sRNA required for function and the length of accessory terminators at the end of constructs to remove unnecessary sequence.

Since the accessory terminators are the largest individual part in the proposed NAR network, we first sought to replace the rrnB terminator originally used in the development of the pT181 attenuator synthetic construct^2^ with a structurally smaller terminator, and then to reduce the repressor sRNA to the minimal structure necessary for function. The rrnB terminator consists of 368 nucleotides and is predicted to fold into a complex structure that could be a source of unwanted RNA-RNA interactions in our NAR network. In previous work we have shown that the rrnB terminator did in fact interfere with the function of a loop-linear chimeric attenuator^32^. Recently, Chen et al.^33^ characterized 582 natural and synthetic transcriptional terminators, which we used as a source for possible terminator alternatives. We chose five terminators that had compact predicted secondary structures and similar or better termination efficiency than that of the rrnB terminator. The terminators were first tested in a simple context to ensure the repressive strength and orthogonality of the repressor RNA were not affected by changing the terminator. To test this, we placed each terminator downstream of the wild type pT181 repressor sRNA, which was constitutively expressed on a high copy plasmid. Each repressor sRNA-terminator construct was tested for repression of the wild type attenuator and its orthogonality to a mutant attenuator *in vivo*, which were placed downstream of the same constitutive promoter and upstream of the super-folder green fluorescent protein (SFGFP) coding sequence^34^ on a medium copy plasmid (Figure S1). Attenuator and repressor sRNA pairs were transformed into *E. coli* TG1 cells, and florescence (485 nm excitation, 520 nm emission) and optical density (OD) (600 nm) were measured for each culture (see Methods). Of the five terminators tested, the repressor sRNA-L3S3P21 terminator construct had the best combination of repression of the wild type attenuator and orthogonality to the mutant attenuator (Figure S1) and was therefore chosen for use in the NAR network.

Next, we tested minimizing the repressor sRNA construct. Early *in vitro* work with the pT181 attenuator showed that the full-length wild type repressor RNA was not required for interaction with and repression of the attenuator^35^. We therefore tested a truncated version of the repressor sRNA, which only contained the hairpin thought to be involved in the initial kissing hairpin interaction with the attenuator. The single hairpin repressor RNA followed by the L3S3P21 terminator showed equivalent repression (90%) of the attenuator when compared to the full-length repressor RNA (Figure 2A).

**Figure 2.**
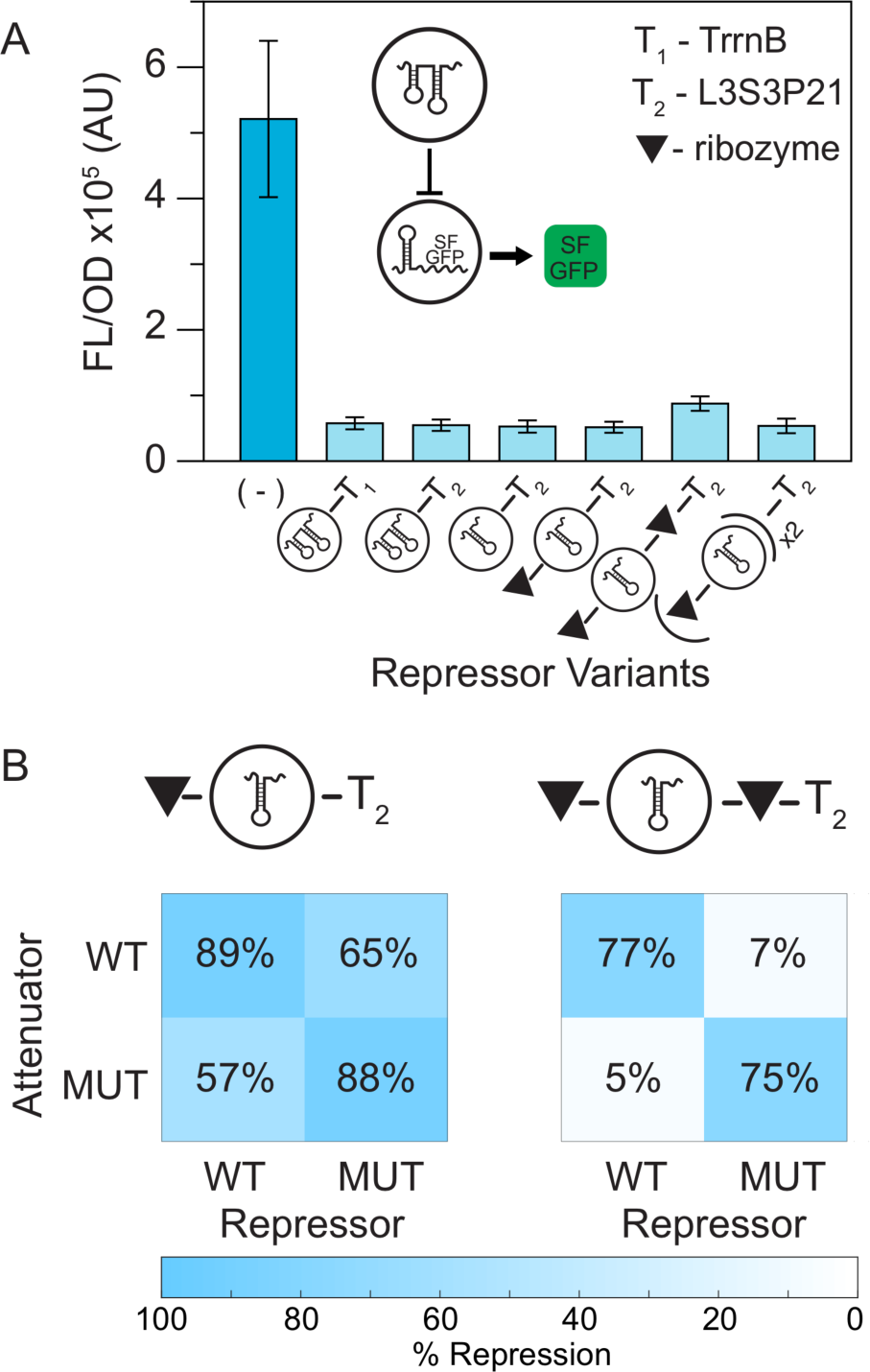
Optimization of RNA parts for network construction. (A) Average steady-state fluorescence (FL/OD) of *E. coli* TG1 cells measuring repression of a construct consisting of the pT181 attenuator controlling SFGFP expression. The pT181-SFGFP construct was transformed with a control plasmid (-), or constructs that expressed variations of the repressor RNA. Variations in the terminator included downstream of the repressor tested the impact of the size of the terminator (T1 vs. T2), with L3S3P21^33^ being smaller than TrrnB. Variations in the size of the repressor RNA repressor (two hairpins vs. one hairpin) tested the impact of including reduced regions of the repressor RNA on repressive function^17^. Other variations tested the impact of using the sTRSV ribozyme for separating RNA parts and including multiple copies of the repressor. Error bars represent standard deviations over nine biological replicates. (B) Orthogonality matrices illustrating that the single hairpin repressor RNA followed by terminator L3S3P21 achieves adequate repression but compromises orthogonality (left). Matrices were generated by measuring the fluorescence of cells transformed with different combinations of the wild type (WT) or specificity mutated (MUT) pT181 attenuator-SFGFP/repressor expression constructs in the repressor configurations indicated by the schematic. Adding a ribozyme in between the single hairpin repressor RNA and terminator L3S3P21 restores the orthogonality (right). Each box in the matrices represents % repression of cells with a repressor RNA expression plasmid compared to cells with the control plasmid. Repression is presented by a color scale in which 100% is blue and 0% is white. Repression % was calculated from the average over nine biological replicates.

In the final NAR construct, it is essential that the individual repressor sRNA copies are free fold into functional conformations and repress the attenuator. It was previously shown that using the hammerhead ribozyme from small tobacco ring spot virus (sTRSV)^36^ to separate copies of repressor RNA on the same transcript improved function of the repressor RNA constructs^2^. Since only the full-length repressor RNA had been tested with the sTRSV ribozyme, we sought to confirm that the sTRSV ribozyme could be used to separate copies of the single hairpin repressor RNA without affecting function. We tested constructs with the sTRSV ribozyme both upstream and downstream of the single hairpin repressor sRNA as well as a double repressor sRNA construct and showed that the sTRSV ribozyme does not interfere with the single hairpin repressor sRNA function (Figure 2A).

Interestingly, while the sTRSV-single hairpin repressor sRNA-L3S3P21 terminator construct showed equivalent repression of the attenuator compared to the construct without the ribozyme; the ribozyme construct was no longer orthogonal to a mutant pT181 attenuator (Figure 2B). However, we observed that adding a second ribozyme downstream of the single hairpin repressor sRNA recovered orthogonality (Figure 2B).

Overall, these optimizations allowed us to remove 387 nucleotides (36% of the total length) from the NAR design. We thus continued constructing and analyzing the sRNA transcriptional NAR with these minimized parts.

### A quantitative model accurately predicts NAR function in TX-TL

Our next goal was to build a more accurate model that could provide quantitative predictions of the NAR network function in TX-TL reactions. To do this, we expanded the two dimensional set of ODE equations that modeled repressor R and reporter MG in Figure 1 to include the maturation steps for both R and MG (Figure 3). In previous work we had found that time trajectories of TX-TL reactions could only be accurately modeled for the pT181 system if a maturation for the repressor sRNA was included^29^.

**Figure 3.**
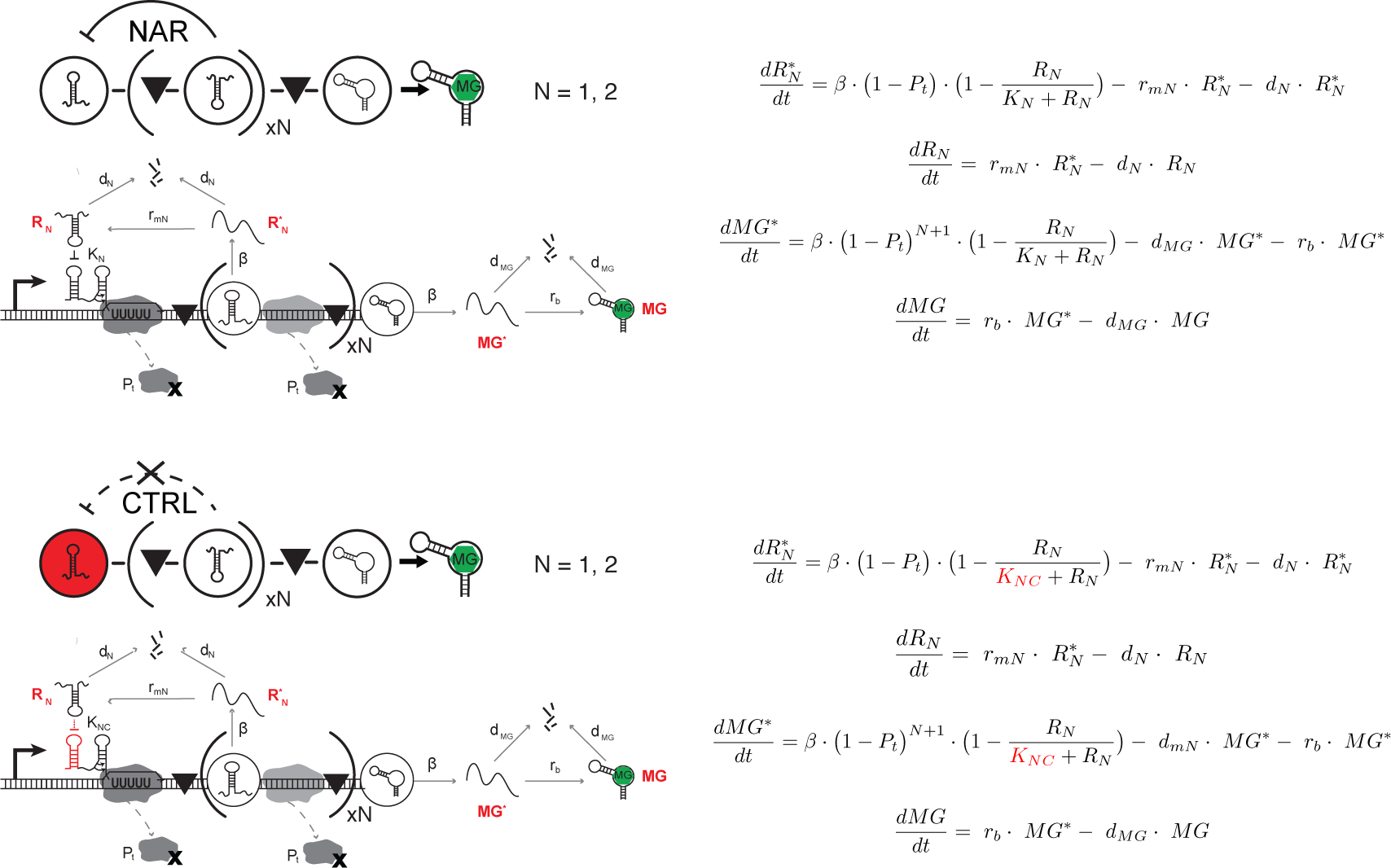
Schematic (left) of the mechanistic steps of the sRNA transcriptional NAR captured by the governing equations (right). Molecular implementations that highlight key rate constants and interactions are shown below each network schematic. N refers to the number of tandem repressor repressors included in each construct. The equations model the tandem copies (N=2) of repressor R_1_ as a single new repressor, R_2_. This effective model also used different degradation rate constants, d_mN_, for each repressor species. Parameter and species descriptions can be found in Table 1 and Table 2, respectively. The only difference between the negative autoregulation (NAR) and control (CTRL) constructs is the use of a mutated repressor RNA that breaks repression, modeled as a change in repression coefficient K_NC_, describing a weak repression caused by crosstalk.

**Table 1.** NAR model parameters.

| Parameter | Description | Unit |
| --- | --- | --- |
| $K_1$ | Repression coefficient of single repressor. | Concentration<br>(Conc.) |
| $K_2$ | Repression coefficient of double repressor. | Conc. |
| $K_{1C}$ | Repression coefficient of single mismatched repressor due to crosstalk. | Conc. |
| $K_{2C}$ | Repression coefficient of double mismatched repressor due to crosstalk. | Conc. |
| $d_1$ | Degradation rate of single repressor. | 1/sec |
| $r_{m1}$ | Maturation rate of single repressor. | 1/sec |
| $r_{m2}$ | Maturation rate of double repressor. | 1/sec |
| $r_b$ | Maturation and binding rate of malachite green (MG) aptamer. | 1/sec |
| $\beta$ | Transcription rate of pre-cleaved MG RNA. | Conc./sec |
| $d_{MG}$ | Degradation rate of MG RNA. | 1/sec |
| $d_2$ | Degradation rate of double repressor. | 1/sec |
| $P_t$ | Probability of transcriptional disruption. | Unitless |

**Table 2:**
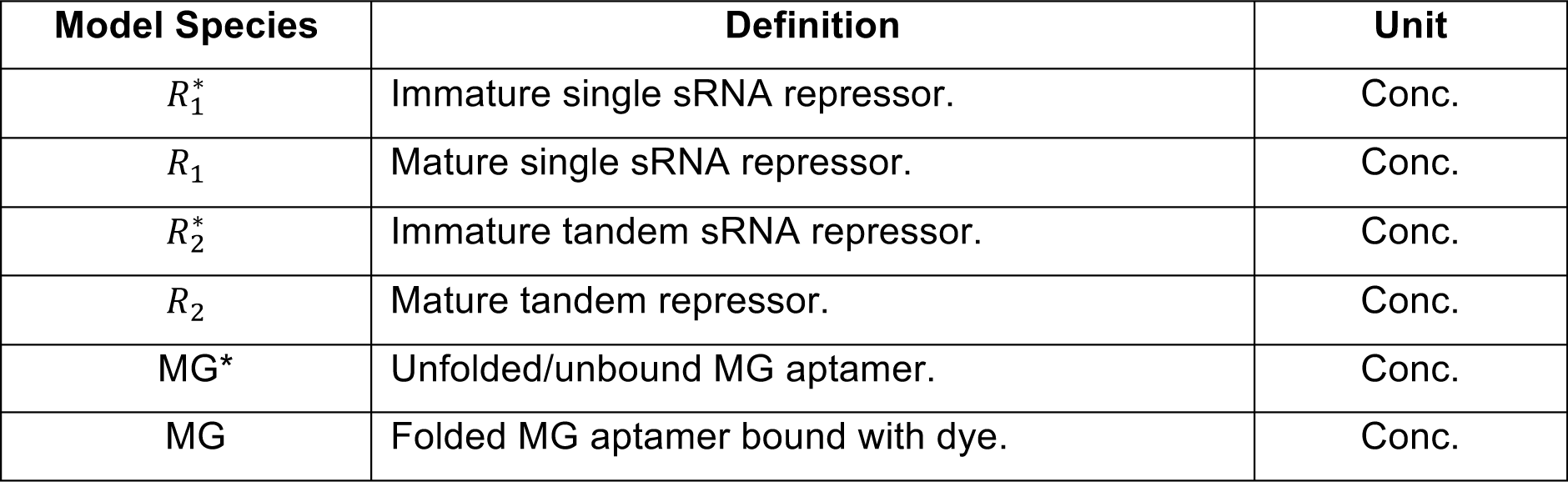
NAR model species.

| Model Species | Definition | Unit |
| --- | --- | --- |
| $R_1^*$ | Immature single sRNA repressor. | Conc. |
| $R_1$ | Mature single sRNA repressor. | Conc. |
| $R_2^*$ | Immature tandem sRNA repressor. | Conc. |
| $R_2$ | Mature tandem repressor. | Conc. |
| $MG^*$ | Unfolded/unbound MG aptamer. | Conc. |
| $MG$ | Folded MG aptamer bound with dye. | Conc. |

In order for the sRNA repressor to function correctly, our previous modeling suggested it has to fold into its functional structure (R) after being initially transcribed as an ‘immature’ species (R*). Similarly, initially transcribed MG molecules (MG*) would need a further step to bind the dye before becoming fully “matured” and observable (MG). Both of these maturation processes were modeled by introducing additional species and specific rates of maturation (Figure 3). For the MG aptamer, we modeled both folding and dye binding with one term, r_b_, to describe the maturation of this RNA reporter. Overall the inclusion of these maturation steps created two additional species to be modeled which expanded the number of ODEs to four.

We also included more detail in our ODE models. Specifically, we introduced terms to account for both autotermination and cross-talk. It is known that there is some amount of autotermination of the pT181 attenuator where transcription stops even in the absence of repressor RNA^2,35^. This autotermination was modeled by including a single parameter that models autotermination probability, P_t_, based on our previous model^29^. In this work, we also found that transcription of repressor sRNA prior to the MG reporter also leads to some small amount of reduction in MG fluorescence (Figure S2A). This indicated that in general the transcription of upstream RNA species reduces the amount of downstream RNA that can be transcribed, likely due to general effects of the imperfect processivity of the RNA polymerase enzyme. To incorporate this in our model, for simplicity we used the same constant parameter P_t_ to denote the probability of transcription being disrupted as polymerase transcribes one of the units of our NAR construct, which could be an attenuator or sRNA-ribozyme construct (Figure 3). Therefore, the probability of a particular sequence region being transcribed without disruption is modeled as (1-P_t_)^n^ where n is the number of construct units upstream (Figure3).

We next incorporated the effect of cross-talk in our model. Experimentally, we sought to characterize the dynamics of the RNA NAR by comparing it to a control construct where only the negative feedback was broken with all other components unchanged. This can be achieved by mutating the attenuator sequence to an orthogonal sequence that does not respond to the repressor RNA^2^ (Figure S1). While in principle this is a good control, the mutant pT181 attenuator is not completely orthogonal to the wild type repressor RNA^2^ (Figure 2B). We therefore had to account for cross-talk in the model in order to accurately model the comparison between the full RNA NAR and the control network. Cross-talk repression for the control network was also also modeled as a first order Hill function with a relatively large repression constant K_c_.

Once our model was proposed, the next step was to determine all 12 unknown parameters (Table 1, Figure 3) that would provide a quantitave estimation of the time-dependent fluorescence trajectories generated by the NAR network in TX-TL. In previous work, we adapted a parameter sensitivity analysis method to guide the design of a series of TX-TL experiments to determine the unknown parameters^29^. For a given experiment modeled by a system of ODEs, sensitivity analysis determines which parameters cause the largest changes to the predicted behavior of the ODEs, i.e. which set of parameters the ODEs are most sensitive to. The most sensitive parameters can be found by computing the sensitivity matrix, which describes how the time varying molecular concentrations of the genetic network in the experiment change in response to a change in the parameters of the model (see Methods). The most sensitive parameters for each experiment are then ideal candidates to extract by fitting the parameters to the experimental data^29^. Since multiple parameters can be estimated from a single experiment, it is therefore possible to estimate all relevant parameters from a reduced set of experiments that are each sensitive to different subsets of parameters.

We set out to perform sensitivity analysis on our NAR network, however because of the feedback element in the NAR, the network is non-linear. Sensitivity analysis of non-linear networks can result in coupled parameter sets^37^ meaning the parameters are restrained by the others within the same set. To avoid solving for coupled parameters, we proposed a set of experiments consisting of key components of the NAR network arranged in configurations that removed network non-linearity (Figure S3). Many of the constructs had already been characterized in previous work with SFGFP as the reporter instead of MG^29,32^. We therefore chose to continue to use SFGFP as the reporter for most of the initial parameterization experiments with one extra experiment to estimate key MG parameters. The use of SFGFP in these experiments however, resulted in four additional parameters to solve for (Figure S3, Table S1). To perform the sensitivity analysis, we used an initial set of 16 parameter guesses taken from our previous work to calculate the sensitivity matrix from our proposed experimental design. We followed the guidelines from our previous study^29^ and designed TX-TL experiments that could be performed at 29°C for 100 min with fluorescence monitored every 5 min. The sensitivity analysis on the proposed linear components from our NAR network suggested that we could identify all 16 parameters with five TX-TL experiments (Figure S3, Supplementary Notes 1 and 2).

We next sought to estimate the parameters in our model by performing the set of experiments designed through the sensitivity analysis. We performed replicate TX-TL experiments with each of the designed plasmid combinations (Figure S3) and collected dynamic fluorescence trajectories of the reactions (Figure S4). Using this experimental data, we then estimated parameters using an iterative fitting procedure that used each experiment in turn to find its designated parameters (see Methods). To capture the natural variation in experimental conditions, we initially input sets of parameters, drawn from uniform distributions centered around an initial best guess, into the fitting procedure and optimized the values of each parameter within each input set (see Methods). This resulted in 10,000 sets of the 16 estimated parameters. To validate our method, we compared measured fluorescence trajectories for each experiment to simulated trajectories using 1,000 sets of randomly drawn parameters from the 10,000 parameter set distributions. This allowed us to compare the experiments to the mean simulated trajectory and 95% confidence intervals, which captured the range of trajectories predicted by our parameter distributions. As shown in Figure S4, each mean simulated trajectory accurately described the experimental observation, with almost every measured experimental trajectory lying within the 95% confidence intervals of our simulation.

Our goal of building a quantitative model was to accurately predict the behavior of the sRNA transcriptional NAR network. To assess the quality of the model prediction, we simulated dynamic trajectories of two hour TX-TL experiments for the four constructs shown in Figure 4A, a no-autoregulation control and NAR network for both the single and double repressor RNA case. This was done using 1,000 sets of randomly drawn parameters from the 10,000 parameter set distributions (Figure 4B-C).

**Figure 4.**
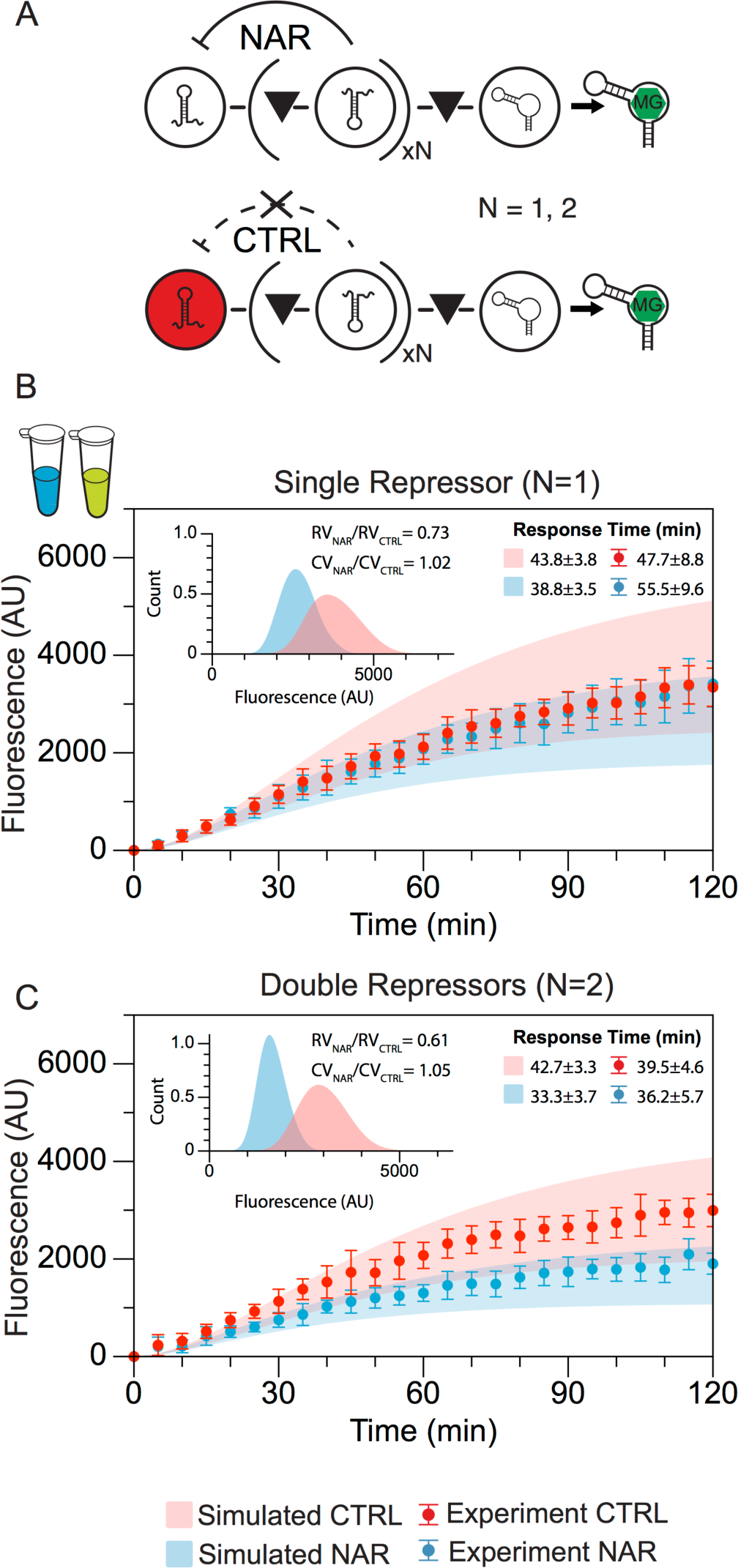
Model prediction and experimental characterization of the sRNA transcriptional NAR constructs in TX-TL. (A) Schematics of the NAR and control constructs designed for TX-TL using malachite green (MG) as a network reporter. (B) and (C) Comparison of experimental trajectories of MG fluorescence in TX-TL experiments (solid circles with error bars) with simulated model predictions (shaded regions) for single repressor NAR (N=1) and double repressor NAR (N=2), respectively. Experimental error bars represent standard deviations of nine individual TX-TL reactions. Model simulated trajectories were generated by performing 1000 simulations with parameters randomly drawn from the set of 10,000 determined from the estimation procedure (see Methods). Predicted and measured response times for all four trajectories are shown in the upper right corner. Inset: The fluorescence at 2 hours for each of the 1000 simulated trajectories are plotted in the histogram and shown in the left upper corner insets. The ratio of relative variance (RV_NAR_/RV_CTRL_) and coefficient of variation (CV_NAR_/CV_CTRL_) indicates that the double repressor NAR construct is also predicted to reduce steady state noise.

To analyze these simulated network predictions, we first took the steady state output of each set of simulated 1000 trajectories and plotted them in a histogram graph to look at the distribution of simulated steady state signal and examine their noise characteristics (Figure 4B-C, Insets). Noise reduction is often measured by coefficient of variance (CV) and relative variance (RV), or “Fano factor”. Because negative autoregulation typically reduces both variance and mean (average) of steady state gene expression, it is important to consider both effects individually^38^. We therefore calculated the RV_NAR_/RV_ctrl_ and CV_NAR_/CV_ctrl_ for both single repressor and double repressor cases. Our model predicted that both versions of the sRNA transcriptional NAR would reduce the relative variance (RV), but not the coefficient of variation (CV). At the same time, the double repressor NAR is predicted to have a stronger RV reduction effect. This finding agreed with previous studies on noise reduction properties of protein mediated transcriptional NARs and sRNA mediated translational NARs^38^, which showed a decrease in RV and increase in CV for both cases. In addition to these noise characteristics, our model predicted that both the single and double repressor NARs would reduce the response time compared to the control network, though with a larger reduction in the case of the double repressor network.

To test our model predictions, we next performed TX-TL experiments with the same constructs to obtain experimental fluorescence trajectories. For all four constructs, experimental results agree with the 95% confidence interval of our simulations (Figure 4B-C). We next sought to compute the response times and noise characteristics of the experimental trajectories. However, there was considerable fluctuations in individual trajectories which precluded noise analysis and made calculating the response time difficult (Figure S5). To overcome this, we generated a smoothened version of experimental trajectories for response time estimation (see Methods, Figure S5). As suggested by the simplified model, the sRNA NAR network with tandem copies of repressor sRNA exhibited a clear separation of fluorescence trajectories with a faster average response time, while the single repressor RNA version did not demonstrate these key characteristics in both simulated and experimental results. We note however that the noisiness of the trajectories caused a large variation in individual trajectory response times making it difficult to quantitatively establish how much faster the double repressor network responded. We did observe that there was a significant separation of trajectories between the NAR and control networks only in the double repressor case.

Overall this result demonstrated that the sRNA transcriptional NAR network functions in cell-free systems and extended the application of our effective model and parameterization method to a non-linear network.

### The sRNA transcriptional NAR network functions *in vivo*

We next sought to adapt the NAR network for use in cells. Since the MG aptamer is difficult to express in cells^31^, to accomplish this we first replaced the MG reporter with a fast-degrading protein reporter, yem-GFP (Figure S6)^39^. Initially, we tested the yem-GFP reporter in a simple context where the expression of yem-GFP was controlled by the pT181 attenuator in *E. coli* cells in the presence or absence of repressor sRNA. We found that the no-repressor GFP expression was lower than that of SFGFP, but the repression with repressor sRNA was nearly 100% (Figure S6A). Next, in order to be able to assess cellular network dynamics from a defined starting time point, we replaced the constitutive promoter used above with the N-acyl homoserine lactone (AHL) inducible promoter P_lux_^40^. We compared the expression of the attenuator-yem-GFP construct under the control of the constitutive promoter and the P_lux_ promoter with varying concentrations of AHL and found that 100 nM of AHL resulted in maximum expression from the promoter (Figure S6B).

Using these two modifications, we built NAR constructs analogous to those in Figure 3A and tested them in *E. coli* cultures while monitoring fluorescence every 20 minutes (Figure 5, S7). As the model and TX-TL results suggested, the NAR construct with a single copy of repressor RNA did not show a significant difference in response time *in vivo* (Figure S8). However, experiments comparing the double repressor RNA NAR network to the control showed that the yem-GFP trajectories began to reach steady state for the NAR network after approximately two hours while the control trajectories were still increasing. To calculate a response time for each construct, we performed regression analysis on data points to determine the steady state of individual fluorescence trajectories. We then used the steady state level to find the time point at which each trajectory reached half of its steady state florescence value (see Methods). The response time (T_1/2_) for the double repressor NAR network was 80 ± 6 min, while the control showed a 97± 5 min response time. This confirmed that the RNA-only NAR network functioned properly *in vivo*.

**Figure 5.**
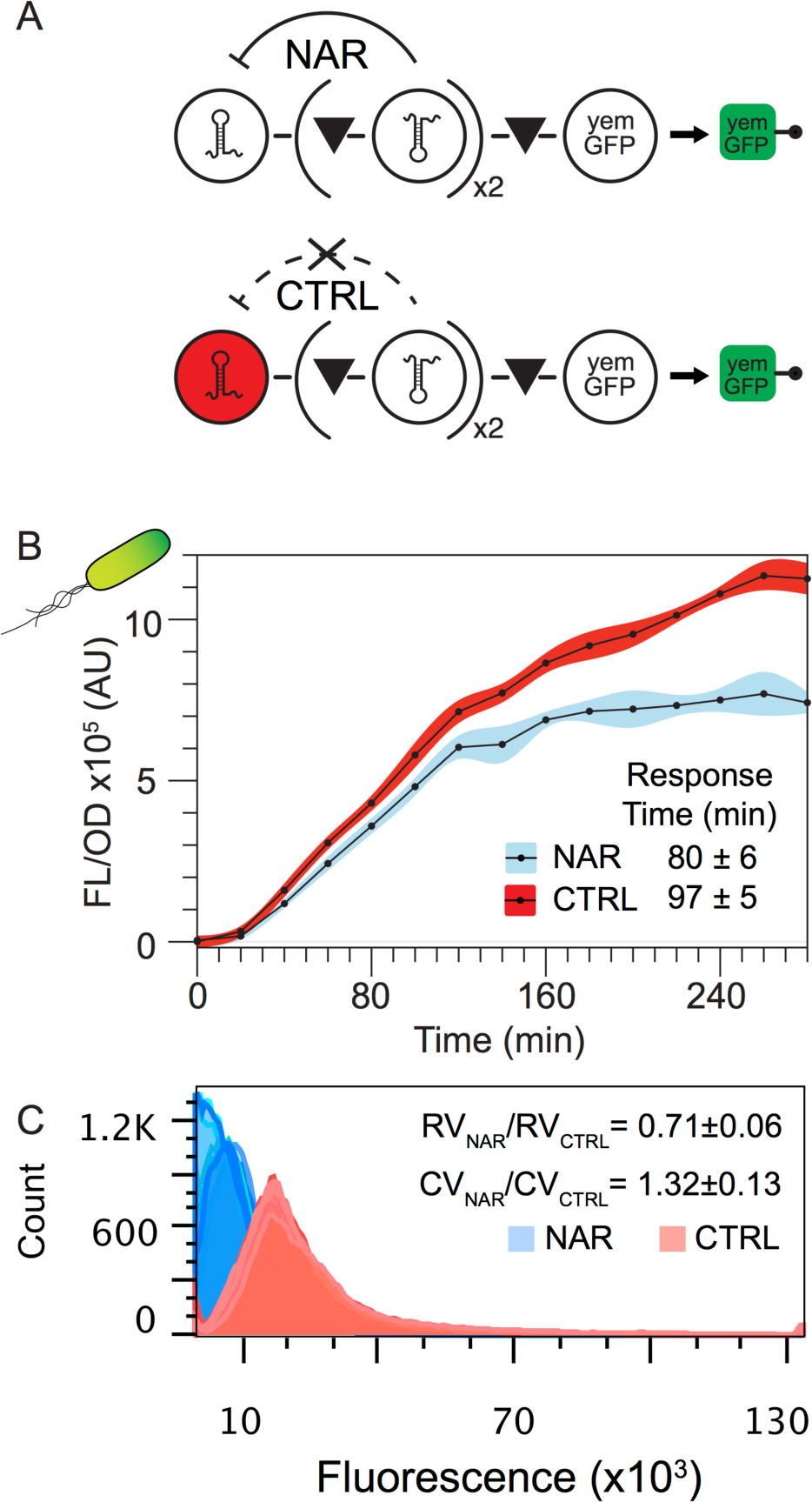
Demonstration and characterization of a functional sRNA transcriptional NAR *in vivo*. (A) Schematics of the NAR and control constructs designed for *in vivo* testing using yem-GFP as a network reporter. (B) Time course of average fluorescence (FL/OD) collected from *E. coli* TG1 cells containing the double repressor NAR (blue) or control (red) construct over a five-hour period. Shaded regions represent standard deviations collected at each time point for nine biological replicates. Calculated response times (see Methods) indicate a functional NAR network that decreases the response time with respect to the control. (C) Flow cytometry histograms of a single time point measurement after 5 hours, tested with the double repressor NAR (blue) and control constructs (red) *in vivo*. Individual distributions are overlaid on each other with different shades of blue and red. The ratio of relative variance (RV) and coefficient of variation (CV) was calculated for each distribution and averaged. The averages of the ratio of (RV_NAR_/RV_CTRL_) and (CV_NAR_/CV_CTRL_) are shown in the upper right hand corner with errors representing the standard deviation of these calculated ratios.

As a further analysis, we also examined the noise reduction effect of sRNA transcriptional NAR networks in *in vivo*. To do this, we performed a flow cytometry analysis of the double repressor NAR and its control for a specific time point at 5 hours after AHL induction (see Methods). We then calculated RV and CV from flow cytometry distributions to characterize network variation. All replicates showed a clear reduction in RV and an increase in CV (Figure 5C). This result is consistent with our simulated noise profile of the sRNA transcriptional NAR in TXTL with estimated parameter distributions. It is also in agreement with previous studies done on protein-mediated transcriptional NARs and sRNA mediated translational NARs, in which both demonstrated decreases in RV and increases in CV^38^.

Overall these results represent the first demonstration of an RNA-only transcriptional NAR network that functions inside living cells.

### Conclusion

In conclusion, we demonstrated the utility of using effective models in combination with TX-TL experiments to systematically and rapidly prototype RNA network designs. Using this design and prototyping method, we were able to create the first functional sRNA mediated transcriptional NAR that functions both in TX-TL and *in vivo*.

We started the design process using a set of over-simplified ODEs to predict potential problems of initial NAR designs and to propose that a design that used a double sRNA repressor would be necessary to observe a reduction in response time that is characteristic of NARs. Using this model-guided design, we also significantly shortened the length of the construct that implemented this design, while preserving the proper functioning of all of the component parts. Similar to a recent investigation^17^, we showed that the truncated version of the sRNA repressor is adequate for achieving the same repression strength of the full-length repressor. However, when a more compact transcriptional terminator is used, this truncated sRNA repressor RNA requires an extra piece of ribozyme between the repressor and the terminator to preserve orthogonality. This result offers an engineering guideline to concatenate multiple RNA components together for more complex sRNA networks.

Once our model had identified the overall NAR design, we then adapted our previously developed sensitivity analysis based parameterization method to construct and parameterize a more detailed model of the network. Using this approach, we designed five simple TX-TL experiments that we used to parameterize all unknown parameters. These parameters were then used to quantitatively predict the dynamic trajectories of two versions of the sRNA transcriptional NAR network. The model prediction suggested that the double repressor version would function according to the two key characteristics of NAR networks: faster response time and lower noise. TX-TL characterization of the network revealed that the model predictions were both qualitatively and quantitatively accurate. Finally, we modified the TX-TL-prototyped NAR constructs for *in vivo* experiments and showed that the network also demonstrated faster response time and lower noise in *E. coli*. Interestingly, the reduction in noise followed a similar trend for our sRNA mediated transcriptional NAR as that observed for protein mediated translational NARs and sRNA mediated translational NARs^38^.

Several aspects of this work are significant. First, this work represents the first transcriptional sRNA negative auto-regulator that functions both in cell free reactions and in living cells. Second, in this work we extended our parameterization procedure to a non-linear network. Because of the natural feedback component of NAR, it is difficult to parameterize its parameters independently. By parameterizing components individually, we were able to incorporate the feedback non-linearity at a later modeling step which we showed was able to recapitulate experimental results. Finally, this work represents a streamlined and quantitative model guided design and TX-TL based prototyping methodology for in-cell RNA network engineering. We found that the parameters we obtained from TX-TL parameterization procedures are adequate to predict the function of our sRNA transcriptional NAR network accurately and provide insights for TX-TL prototyping experiments that then inform the construction of networks that function in cells. We anticipate this design-build-prototype-test network building process will minimize cloning effort and speed up network development cycles as the field moves towards more sophisticated synthetic network engineering.

## Methods

### Plasmid construction and purification

A table of all the plasmids used in this study can be found in Supporting Information Table S5, with key sequences found in Table S4. The pT181 attenuator and repressor plasmids, pT181 mutant attenuator and repressor plasmids were constructs pAPA1272, pAPA1256, pAPA1273, and pAPA1257 respectively, from Lucks et al.^2^. All NAR and control constructs were created using Golden Gate^41^ assembly and Gibson assembly. The NAR constructs were created with pT181 attenuator and pT181 repressor, the control constructs where created with pT181 mutant attenuator and pT181 repressor. Plasmids were purified using a Qiagen QIAfilter Plasmid Midi Kit (Catalog number: 12243) followed by isopropanol precipitation and eluted with double distilled water.

### TX-TL experiments

TX-TL buffer and extract was prepared according to Garamella et al. (Garamella 2016). TX-TL buffer and extract tubes were thawed on ice for approximately 20 min prior to performing experiments. Separate reaction tubes were prepared with combinations of DNA representing a given network condition. Appropriate volumes of DNA, buffer, and extract were calculated using a custom spreadsheet developed by Sun et al^42^. Buffer and extract were mixed together and incubated for 20 min at 37°C. DNA for the specific experiment was then added into each tube according to the previously published protocol^27^. Ten μL of each TX-TL reaction mixture was transferred to a 384-well plate (Nunc 142761), covered with a plate seal (Nunc 232701), and placed on a Biotek SynergyH1m plate reader. Temperature was controlled at 29°C. Superfolder GFP (SFGFP) (485 nm excitation, 520 nm emission, 100 gain) or malachite green (MG) (610 nm excitation, 650 nm emission, 150 gain) fluorescence was measured every 5 min for 120 min. MG constructs were observed to immediately generate observable signal in between mixing and loading the plate reader, making quantification of early time points difficult. Therefore, the first measurement of all constructs with MG were assumed to be 0 and data collection was timed to start 5 minutes after reactions were mixed. Each reaction was run with a minimum of triplicate technical repeats, and repeated three times independently (minimum of nine total replicates). A ten μL sample of TX-TL buffer, extract, and water was run together with each independent experiment as blank. All data points were then processed with blank values subtracted at each time point.

### Network/Model Simulations

All models were simulated by solving corresponding ODEs using *Matlab* function *ode15s* over a set of discrete time steps using guessed or estimated parameters. All initial conditions for species concentrations were zero.

### Parameterization experiment design

All unknown parameters were estimated by fitting model simulations to a small set of time course data generated from TX-TL experiments. Following previous work^29^, these experiments were designed through an iterative process of identifiability tests on the set of model equations (Figure S3, Supplementary Note 1). An individual experiment was expected to produce a measurable trajectory of fluorescence as a function of time. After a specific experiment was proposed, we determined which parameters were “identifiable” from this experiment through sensitivity analysis (See Supplementary Note 2). The iterative experiment starts with the simplest experiment that has the smallest number of network components and thus the smallest number of parameters governing the fluorescence trajectory. Identifiable parameters from this experiment are then estimated by fitting models to the experimental data as described below (Parameter estimation). The next experiment that contains a greater number of components and the analysis repeated, except that parameters already identified by previous experiments were marked as “determined” and were not included in sensitivity analysis. Rounds of experimental design and sensitivity analysis were performed until all 16 parameters could be identified by 5 TX-TL experiments (Figure S3).

### Parameter estimation

Parameters were estimated from each designed experiment by fitting the identifiable parameters of that experiment to measured fluorescence trajectories. The parameter estimation problem is given by:

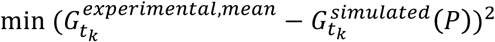

where 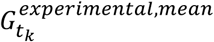 denotes the average value of the experimentally observed SFGFP expression at a certain time t_k_. The vector P contains all of the identifiable parameters in the experiment being analyzed. 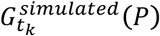 denotes the model simulated SFGFP expression at time t_k_.

To optimize the initial guesses, we first started with a set of initial guess based on pervious work (Table S1)^29^. Then we constructed uniform distributions (100 points) around each parameters value (±15%), generating 100 sets of parameters, which led to 100 simulated dynamic trajectories for each of the 5 estimation experiments designed by sensitivity analysis procedure. We picked the set of parameters that generated the best fit to the 5 sets of experimental data collectively, and used it as a starting point for the next round. We repeated this procedure 10 times to achieve the best guess.

Then we started the estimation process with the best guessed parameter set and constructed uniform distributions (10000 points) around each parameter value (±15%). Each set of parameters served as a different starting point for finding optimal parameter estimates. Each estimate set was found by sequentially applying the fitting procedure to each of the designed parameters using the *Matlab* function *fmincons*, only fitting the identifiable parameters for that experiment. Fit parameters from one experiment were then used to update and replace the initial guesses before moving on to the next experiment until each of the 16 unknown parameters were fit from the 5 experiments. The result of this procedure produces 10000 sets of estimated parameters, which were then subject to the analysis described in the main text.

### Strains, Growth Media and *in vivo* Gene Expression Experiments

All experiments were performed in *E. coli* strain TG1. Plasmids with different network constructs were transformed into TG1 competent cells, plated on LB + Agar plates containing 100 μg/ml carbenicillin, and incubated overnight at 37°C. At least three colonies of each experimental condition was inoculated into 300μL of LB containing carbenicillin in a 2mL 96-well block (Costar 3960), and grown for 12 hours overnight at 37°C at 1000 rpm in a Labnet Vortemp 56 bench top shaker. Four microliters of the overnight culture were added to 196 μl of M9 supplemented M9 media (1X M9 minimal salts, 1 mM thiamine hydrochloride, 0.4% glycerol, 0.2% casamino acids, 2 mM MgSO4, 0.1 mM CaCl2) containing carbenicillin and grown for 4 hours at 37°C at 1000 rpm.

For time course measurements, the sub-culture was then diluted to an 0.015 OD with M9 + carbenicillin and 100nM N-Acyl homoserine lactone (AHLs) to a total volume of 1mL and grown at 37°C at 1000 rpm. During the 5-hour time course, 50μL of culture was removed from of the main culture every 20 minutes for measurements.

Fluorescence (485 nm excitation, 520 nm emission) and optical density (OD, 600 nm) were measured on a Biotek SynergyH1m plate reader.

For steady state expression measurement, the sub-culture was then diluted to an 0.015 OD with M9 + carbenicillin and 100nM *N*-Acyl homoserine lactones (AHLs) to a total volume of 200μl and grown at 37°C at 1000 rpm for 5 hours. Fluorescence (485 nm excitation, 520 nm emission) and optical density (OD, 600 nm) were then measured on a Biotek SynergyH1m plate reader.

For flow cytometry measurement, the sub-culture was then diluted to an 0.015 OD with M9 + carbenicillin and 100nM *N*-Acyl homoserine lactones (AHLs) to a total volume of 3ml and grown in culture tubes for 5 hours at 37°C at 220rpm. A total of 50,000 events were collected on a BD Accuri^TM^ C6 plus Flow Cytometer during measurement.

All experiments contained three replicates from at least three different transformed colonies, and were repeated independently three times for a minimum of nine total replicates.

### Response-time Calculation

To estimate the response time *T*_1/2_ of a network, we estimated the time for individual dynamic trajectories to reach steady state based on the rate production of fluorescence signal. We first scaled all data points by the maximum experimentally observed fluorescence value across all four experiments 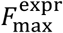:

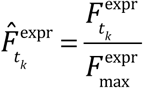

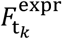 is the experimentally observed fluorescence/OD value (malachite green for *in vitro* networks and yem-GFP for *in vivo* networks) at any time *t_k_* during the experiment. For *in vitro* data sets where the oscillations in fluorescence values could complicate data analysis, we employed the *smooth* function in MATLAB to filter out oscillations in data points. This function smooths the data using a moving average filter with a span of 5 data points (Figure S5, S7, S8).

The least square method was then used to estimate the rate of SFGFP signal production. Beginning with the first normalized data point (*k* = 1) in the data set, we took a total of five data points (*n* = 5) and calculated the least squares slope (*m*) at time point k with following equation:

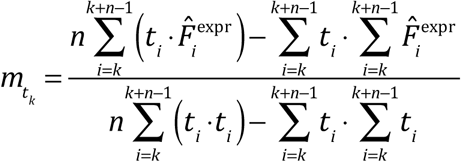

The objective of the least square is to find slope (*m*) that minimizes the sum of squared errors (*r*^2^) of the differences between the observed SFGFP production and those predicted by a linear model. The *m* found became the slope at time point *t*_k=1_, and we step into the next set of five data points with *k* = 2, 3 etc., creating an array of slopes to describe the fluorescence trajectory.

When the absolute value of the calculated slope is below a set threshold of 0.005, the trajectory is defined to be at steady state and its fluorescence level at steady state is recorded. For trajectories that did not reach steady state at the end of the experiment, we forecasted its trajectories beyond the experimental time scope and predicted the time of steady state, assuming the SFGFP signal production rate declines at a constant rate.

Using the steady state signal values, we traced through the trajectory to identify the response time when the network signal reached half of its steady state value. If the exact value was in between two experimental data points, a linear interpolation on the two experimental data points gives an estimate for the response time corresponding to the half of the steady state fluorescence value.

### Signal Noise Analysis

Noise magnitude was considered with two measures: the coefficient of variation (CV) of signal X and the relative variance (RV) or ‘Fano factor’ of signal X. They were calculated with following formulas:

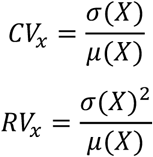

Where *σ(X)* is the standard deviation of signal X and *μ(X)* is the average value of signal X. Mathematically, CV of signal X can increase when the mean decreases while RV does not decrease sufficiently^38^.

## Acknowledgements

The authors thank Marshall Colville for early help in characterizing NAR circuitry, and Adam Silverman for helpful comments on this manuscript. This work was supported by an NSF CAREER Award (1452441 to J. B. L.), a National Science Foundation Graduate Research Fellowship (grant no. DGE-1144153 to M.K.T.) and Searle Funds at The Chicago Community Trust (to J.B.L.).

